# Development of fluoro-7-aminocarboxycoumarin-based mitochondrial pyruvate carrier inhibitors as anticancer agents

**DOI:** 10.1101/2024.05.22.595353

**Authors:** Tanner J. Schumacher, Zachary S. Gardner, Jon Rumbley, Conor T. Ronayne, Venkatram R. Mereddy

## Abstract

Reprogrammed metabolism of cancer cells offers a unique target for pharmacological intervention. In the current study, a series of novel and potentially metabolically stable fluoro-substituted aminocarboxycoumarin derivatives are evaluated for their mitochondrial pyruvate carrier (MPC) inhibition properties. Our studies indicate that the aminocarboxycoumarin template elicits potent MPC inhibitory characteristics, and specifically, structure activity relationship studies show that the *N-*methyl-*N-*benzyl structural template provides the optimal inhibitory capacity. Further respiratory experiments demonstrate that candidate compounds specifically inhibit pyruvate driven respiration without substantially affecting other metabolic fuels consistent with MPC inhibition. Further, computational homology and inhibitor docking studies illustrate that aminocarboxycoumarin binding characteristics are indicative of reversible covalent bonding with amino acids in the pyruvate binding domain. Epifluorescent microscopy experiments illustrated that FACC2 accumulates in the mitochondria to a similar extent as parent 7ACC2. Additionally, lead candidate aminocarboxycoumarin derivative D7 elicits cancer cell proliferation inhibition specifically in monocarboxylate transporter 1 (MCT1) expressing 4T1, consistent with its ability to accumulate intracellular lactate. *In vivo* tumor growth studies illustrate that D7 significantly reduces the tumor burden in two isogeneic murine cell lines 4T1 and 67nr. These studies provide novel MPC inhibitors with potential for anticancer applications.

## INTRODUCTION

The onset of neoplasia results from numerous genetic mutations leading to uncontrolled growth, division and proliferation. Hanahan and Weinberg have defined several classical hallmarks of cancer including evasion of apoptosis, self-sufficiency of growth signals, and insensitivity to anti-growth signals [1]. Further, cancer cells adapt the ability to stimulate the growth of new blood vessels (angiogenesis) and evolve invasion and metastatic aptitude. In this regard, cancer cells exhibit limitless replicative potential forming genomically unstable cellular masses recognized as tumors [1]. Within the tumor microenvironment, unequal distribution of oxygen and nutrients leads to diverse tissue types made up of highly heterogeneous cells where the metabolic and proliferative capability of the tumor depends largely on the extent of hypoxia [1–5]. Oncogenic mutations that initiate the formation of the tumor mass lead to cells that are genomically unstable resulting in the replicative immortality and expansion of the malignancy [2,6].

To expand on the classical hallmarks, numerous enabling characteristics and emerging hallmarks of cancer that support such replicative immortality within the diverse tumor microenvironment have been described [8]. One of the important emerging hallmarks includes the cancer cells ability to deregulate their energetics resulting in reprogrammed metabolism [5]. To keep up with the large energy requirement of neoplastic proliferation, cancer cells exhibit marked metabolic shifts from what is observed in normal quiescent epithelial tissue [2,5,7–9]. In non-malignant cells, energy is obtained via an efficient mitochondrial oxidative phosphorylation (OxPhos), where one mole of glucose yields 36 moles of ATP. To do so, normal cells transfer high energy electrons from NADH from several other biosynthetic pathways into the electron transport chain (ETC). During which, pyruvate enters the tricarboxylic acid (TCA) cycle through many coupled enzymatic redox reactions which together reduces molecular oxygen to two moles of water. Warburg observed that cancer cells largely undergo a metabolic switch from OxPhos to the more energy inefficient glycolysis, even under sufficient oxygen conditions [7,10–12]. To keep up with the high energy demand of cancer progression, glycolytic related enzymes and transporters including glucose transporters (GLUT1), hexokinase (HK), glucose 6 phosphate dehydrogenase, monocarboxylate transporters (MCTs) and others have been shown to be highly over-expressed in several malignancies [7]. This aerobic glycolysis, termed Warburg effect, has been observed in numerous cancer types under both oxygen poor and rich conditions, and has emerged as a defining characteristic of cancer cell metabolism [7,10–12].

Recent evidence in contrast to the Warburg effect postulates that cancer associated stromal fibroblasts are stimulated to upregulate glycolysis, and the metabolic byproducts namely lactate and pyruvate, are shuttled from the stromal compartment to the cancer epithelial cells for mitochondrial OxPhos [13–15]. This shift not only increases energy production, but also enables metabolite cycling through the TCA cycle where metabolic intermediates are important for biomass generation [13–15]. This phenomenon has been aptly termed the reverse Warburg effect, and demonstrates the unique ability of tumors to recruit non-malignant tissues and undergo metabolic symbiosis to obtain energy and biosynthetic starting materials in support of rapid and uncontrolled cell division.

One of the important molecular players in coupling glycolysis and mitochondrial OxPhos is the mitochondrial pyruvate carrier (MPC). The MPC facilitates the transport of cytosolic pyruvate, either as a byproduct of glycolytic metabolism or influxed by monocarboxylate transporters, into the mitochondrial matrix [16,17]. The MPC1 and MPC2 genes encode two obligate protein subunits of the MPC that form a heteroligomeric complex wherein both proteins are required for activity as loss of one leads to destabilization and degradation of the MPC complex [17]. The MPC is found on the inner mitochondrial membrane and imports the metabolic end product of glycolysis, pyruvate, into the mitochondrial matrix for incorporation into intermediary metabolism in the citric acid cycle (TCA) [16,17]. Thus, MPC couples the two major energetic pathways, glycolysis and OxPhos, for energetic and biosynthetic needs of the rapidly proliferating cancer cells. Importantly, recent evidence suggests that highly oxidative cancer cell types exhibit increased levels of mitochondrial respiration and anabolic processes that drive cancer cell proliferation [18]. Hence, targeting of MPC has high therapeutic potential for the treatment of a wide variety of cancers that depend on metabolic plasticity.

We and others have recently developed new generation *N,N-*dialkylcyanocinnamic acid and aminocarboxycoumarin derivatives with potent inhibition properties of MCT1 and MCT4 mediated lactate influx [19–22]. Recent evidence suggests that inhibition of mitochondrial pyruvate flux leads to increased cytosolic pyruvate concentrations that can modulate MCT mediated lactate uptake [18]. In this regard, we reason that our first-generation inhibitors may be acting on the MPC resulting in feedback mediated inhibition of MCTs as we previously reported. In the current study, we have synthesized and evaluated a series of novel aminocarboxycoumarin-based MPC inhibitors for the treatment of cancer.

## RESULTS AND DISCUSSION

### Design and synthesis of fluoro-aminocarboxycoumarin (FACC)-based mitochondrial pyruvate carrier inhibitors

We have reported on the synthesis and evaluation of novel N,N-dialkylcyanocinnamic acid and carboxycoumarin derivatives as inhibitors of MCT1 and MCT4 mediated lactate uptake for anticancer applications. Recently, it has been demonstrated that the ability of candidate compounds to inhibit lactate influx was due to MPC inhibition leading to intracellular pyruvate levels that ultimately feedback-regulate MCT1/4 facilitated lactate flux. In this regard, lead aminocarboxycoumarin inhibitor 7ACC2 has been evaluated for its anticancer properties (Figure 1). Similarly, we have identified a lead candidate based on the *N,N*-dialkylcyanocinnamic acid (2CAA) series that potently inhibits lactate influx and exhibits significant tumor growth inhibition properties as a single agent. However, we have found that these compounds exhibit very limited metabolic stability with low biological half-lives and rapid clearance rates. We attribute these pharmaceutical shortcomings to un-substituted phenyl and benzyl groups on 2CAA and 7ACC2, respectively, for rapid CYP450 mediated hydroxylation (**Scheme 1A**). Further, the *N-*benzyl and *N-*methyl carbons on 7ACC2 may become susceptible to CYP450 oxidative cleavage. Hence, we sought to synthetically modify 7ACC2 with fluoro-substituted phenyl and benzyl groups to aim for maintained pharmacological potency and improved pharmaceutical properties, namely improved biological half-life (FACC1, FACC2, FACC3, **Scheme 1B-C**).

**Figure 1.**
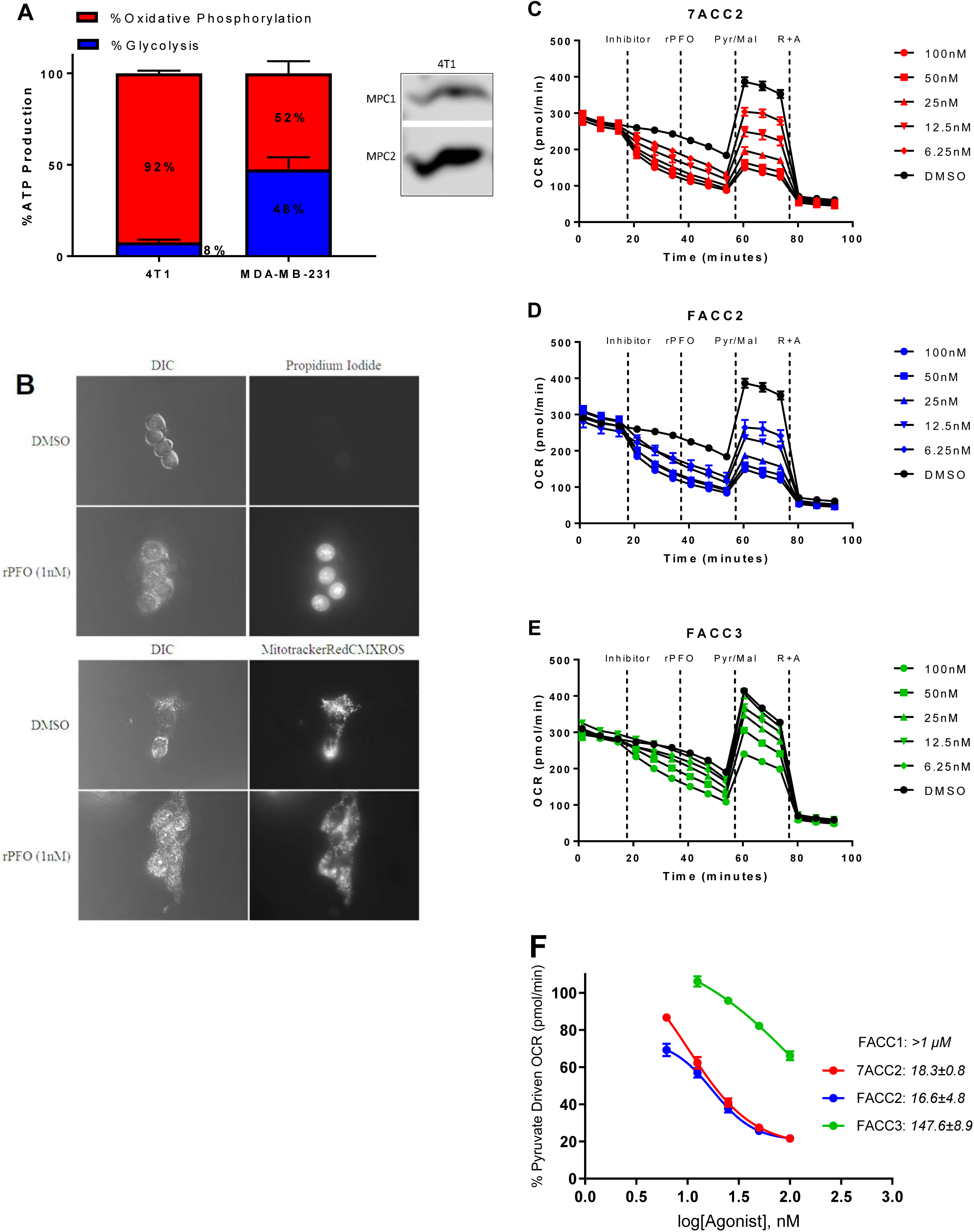
Candidate compounds inhibit pyruvate driven respiration in (**A**) highly oxygen consuming and MPC1/2 expressing 4T1 cells. (**B**) Microscopy experiments illustrate that rPFO (1nM) effectively permeabilized 4T1 cells as indicated by propidium iodide uptake without altering mitotracker red fluorescence. (**C-E**) Candidate compounds inhibit pyruvate driven respiration in permeabilized cells in a dose dependent fashion enabling (**F**) dose response curves to be generated and IC_50_ values to be calculated. All data are representative of at least three independent experiments.

**Scheme 1.**
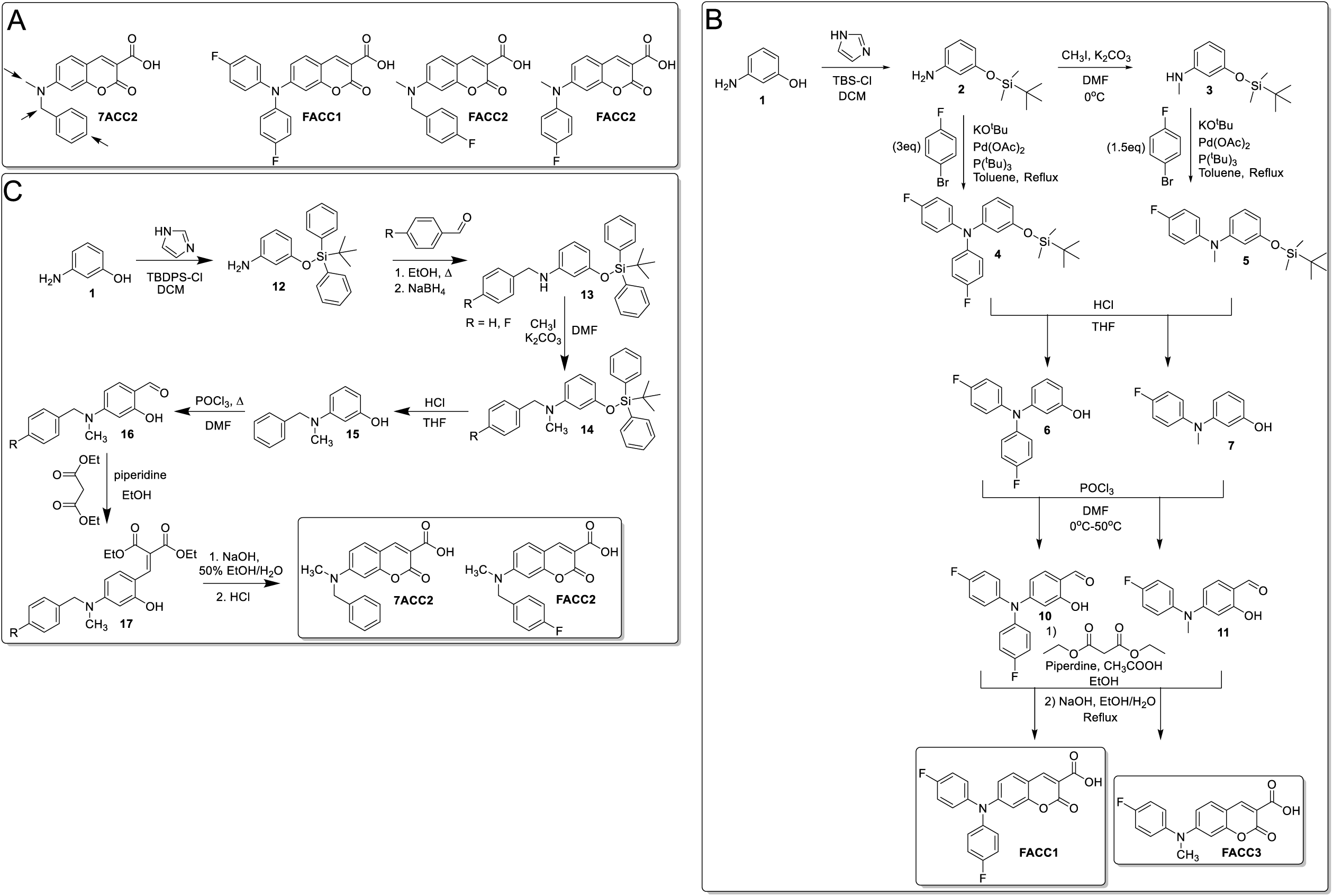
Synthesis of FACC1, FACC2, 7ACC2, and FACC3.

To synthesize the corresponding FACC derivatives, the hydroxyl group of starting material 3-aminophenol **1** was first protected with ^t^butyldimethylsilyl chloride (TBS-Cl) to obtain the corresponding silyl protected amino phenol **2** (**Scheme 1B**). At this stage, **2** was utilized in the synthetic scheme of FACC1 and FACC3 with varying synthetic protocols. Towards the synthesis of FACC1, **2** underwent selective nucleophilic aromatic substitution onto *p*-fluoro-bromobenzene under catalytic Pd(OAc)_2_/P(^t^Bu)_3_ conditions in the presence of KO^t^Bu (**Scheme 1B**). The resulting bis-fluorophenyl-o-silyl protected **4** was further deprotected under acidic conditions to provide the corresponding 3(bis(fluorophenyl)amino)phenol **6**. Direct Vilsmeier-Haack formylation of **6** resulted in the corresponding 4-(bis(4-fluorophenyl)amino)-2-hydroxybenzaldehyde **10**. Likewise, **10** was condensed with diethyl malonate under Knoevenagel conditions in the presence of piperidine and acetic acid in methanol. Subsequent hydrolysis using sodium hydroxide resulted in the product FACC1 (**Scheme 1B**).

The synthesis of FACC3 was initiated with selective mono-methylation of **2** with methyl iodide to obtain 3-((tert-butyldimethylsilyl)oxy)-N-methylaniline **3** (Scheme 1). Selective nucleophilic aromatic substitution of **3** onto *p*-fluoro-bromobenzene under catalytic Pd(OAc)_2_/P(^t^Bu)_3_ conditions in the presence of KO^t^Bu was again performed resulting in the 3-((tert-butyldimethylsilyl)oxy)-N-(4-fluorophenyl)-N-methylaniline **5** (Scheme 1). Acid-mediated deprotection of **5**, followed by Vilsmeier-Haack formylation of **7** provided the respective 4-((4-fluorophenyl)(methyl)amino)-2-hydroxybenzaldehyde **11**. In parallel fashion, Knoevenagel condensation of **11** with diethyl malonate in the presence of piperidine and acetic acid yielded the product FACC3 (**Scheme 1B**).

A slightly modified scheme was utilized to obtain 7ACC2 and FACC2. Initial silyl protection of 3-aminophenol **1** was performed using ^t^butyldiphenylsilyl chloride (TBDPS-Cl) to provide the corresponding o-protected aminophenol **12** (Scheme 2). Reductive amination of **12** with either benzaldehyde or p-fluorobenzaldehyde resulted in the corresponding benzyl or fluoro-benzyl **13.** Methylation utilizing methyl iodide under basic conditions provided the equivalent *N*-methyl **14**, which was further deprotected under acidic conditions providing the 3-(benzyl(methyl)amino)phenol **15** (Scheme 2). Above mentioned Vilsmeier-Haack formylation conditions provided the corresponding aldehyde **16**. Knoevenagel condensation of **16** with diethyl malonate, followed by hydrolysis provided the 7ACC2 and FACC2, respectively (Scheme 2). It is noted that earlier generation compound 2CAA was synthesized using procedures reported elsewhere.

**Scheme 2.**
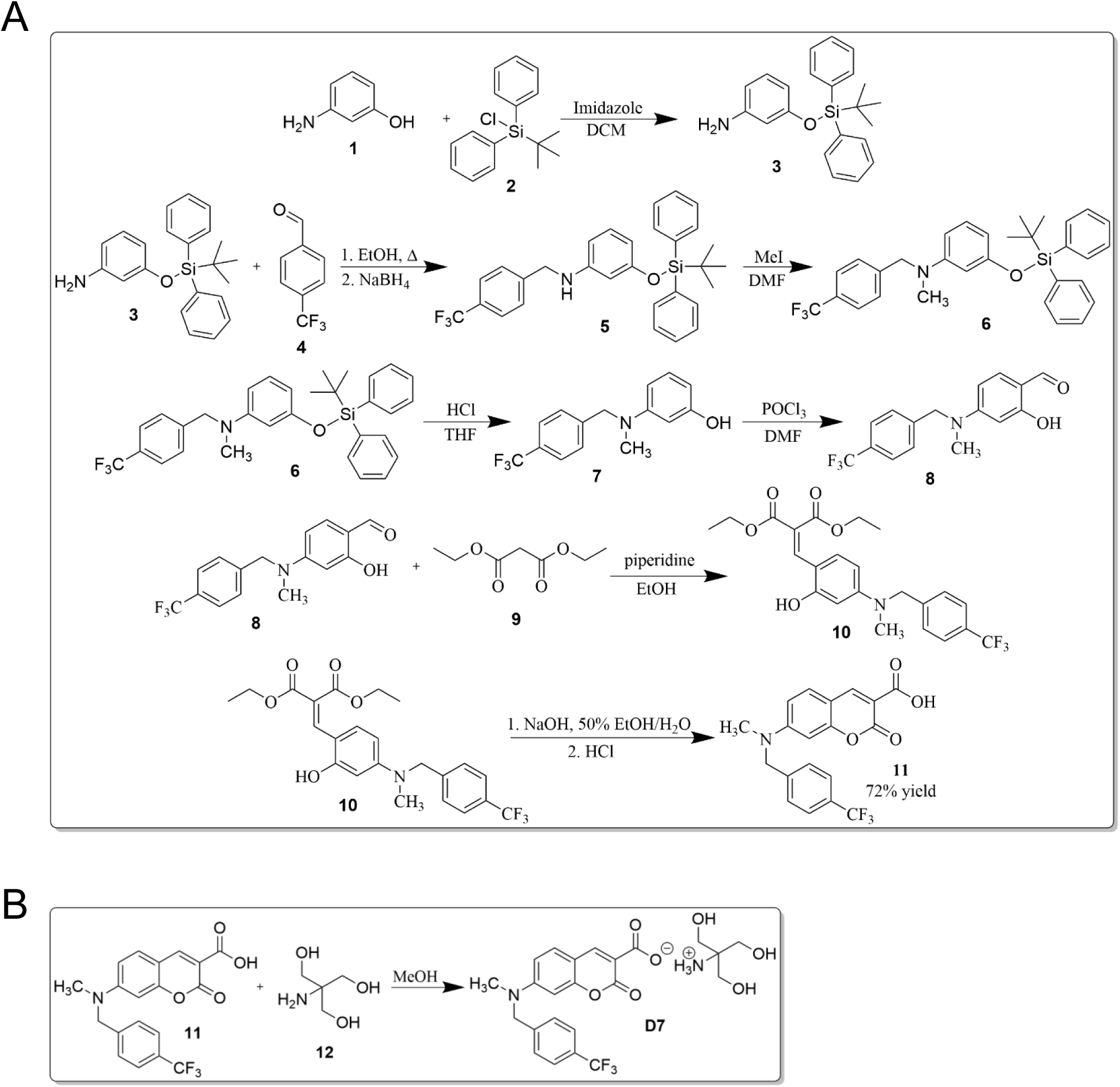
Synthesis of D7.

### FCAA and FACC drug candidates exhibit potent inhibition properties of pyruvate driven respiration in 4T1 cells

To validate that the FACC inhibitors retained the pharmacological properties of 7ACC2 we evaluated their efficacy to inhibit pyruvate driven respiration in 4T1 breast cancer cell line. This cell line was found to exhibit a substantially high basal level of oxidative phosphorylation when compared to an established glycolytic cell line, MDA-MB-231 (**Figure 1A**). Further, this cell line was found to express both MPC1 and 2 at readily detectable levels, suitable for screening drug candidates with potential MPC inhibitory capacity (**Figure 1A**). To evaluate the ability of candidate compounds to inhibit pyruvate driven respiration (MPC inhibition), we employed a slightly modified Seahorse XFe96 based respiratory experiment in permeabilized cells that has been utilized elsewhere. In permeabilized cells, small polar metabolites, including pyruvate, can be delivered directly to mitochondria independent of plasma membrane transport and hence, the kinetics of mitochondrial pyruvate driven respiration can be measured directly. Recombinant perfringolsyin O (rPFO) is a cytolysin excreted by *Clostridium perfringins* that potently and acutely permeabilizes the plasma membrane of eukaryotic cells without impacting organelle membrane integrity. To test the ability of rPFO to permeabilized the plasma membrane in our model system, we employed epifluorescent microscopy experiments (**Figure 1B**). Here, 4T1 cells were seeded in MatTek glass-bottom dishes and were incubated for 48 hours for adherence. Cells were than exposed to rPFO (1nM) and incubated for 30min in growth media, a time point relevant to the exposure in Seahorse experiments. To validate membrane permeability, propidium iodide uptake was observed. Media was then aspirated and was replaced with a mannitol/sucrose buffer (MAS buffer; 70mM sucrose, 220mM mannitol, 10mM potassium phosphate monobasic, 5mM magnesium chloride, 2mM HEPES, and 1mM EGTA) containing both propidium iodide (PI) and mitotracker red CMXROS (MTR) – a probe that binds and accumulates to the mitochondria as a function of membrane potential. PI enabled observation of membrane permeability as intact membranes do not allow PI uptake and hence. MTR was utilized in these experiments to assess the effects of rPFO on mitochondrial viability as damaged mitochondria will exhibit a dim/diffuse fluorescence. These experiments revealed that rPFO potently (1nM) and acutely (30min) permeabilized the plasma membranes as indicated by PI uptake in rPFO treated cells (**Figure 1B**). Further, we observed that mitochondria remained viable with comparable MTR fluorescent intensity in rPFO cells when compared to the controls (**Figure 1B**).

After validating the ability of rPFO to permeabilize 4T1 cells without damaging mitochondria, we sought to investigate the ability of candidate compounds to inhibit pyruvate driven respiration in these cells (**Figure 1C-F**). In this regard, 4T1 cells were seeded in Seahorse XFe96 well plates and were incubated to reach confluence (∼18-24 hours) in growth media (+serum). Growth media was then aspirated and cells were washed with MAS buffer to remove serum and endogenous metabolic substrates. Serum- and substrate-starved cells were then incubated for equilibration in MAS buffer in a non CO_2_ incubator. Inhibitors, rPFO, and substrate milieus were prepared in MAS buffer to be injected in succession to allow for real-time observation of the effects of compound-treated cultures on oxygen consumption rates (OCR) when compared to vehicle (DMSO) treated cells. Equilibrated in-tact cells were allowed to establish basal respiratory rates, followed by the injection of candidate compounds 7ACC2, FACC1, FACC2, and FACC3 at varying dose-titrated concentrations, respectively, and acute decreases in OCR were observed (**Figure 1C-F**). Compound exposure was performed in intact cells to avoid influence of rPFO on acute cellular targets. Cells were then exposed to rPFO, followed by the FCCP stimulated pyruvate substrate milieu which consisted of pyruvate (Pyr), malate (Mal), and dichloroacetate (DCA) to fuel uninhibited pyruvate respiratory flux and maximal respiration. Malate and DCA were included to allow for continuous TCA cycle function without acetyl CoA feedback mediated inhibition (malate) or pyruvate dehydrogenase regulated inhibition (DCA) of pyruvate uptake. Here, we observed a dose-dependent inhibition of pyruvate driven respiration in cells treated with the candidate compounds, allowing for logarithmic dose-response curves to be generated, and 50% inhibitory concentrations (IC_50_) to be calculated (**Figure 1C-F**). Finally, complex I and II inhibitors rotenone and antimycin A were injected to halt respiration and end the assay. These studies revealed that candidate inhibitors exhibited a range of IC_50_ values and gave important insights into structure activity relationships. Interestingly, the bis-fluorophenyl FACC1 completely lost activity when compared to parent 7ACC2 as inhibitory concentration was not reached below 1µM when compared to 18nM of 7ACC2 (**Figure 1F**). Fluoro-substitution of FACC2 did not alter the IC_50_ when of the parent with equipotent 16nM inhibitory capacity (**Figure 1F**). Interestingly, removal of the benzylic carbon in the *N-*methyl-*N-*fluorophenyl example resulted in decreased activity with an IC_50_ value of 147nM, indicating a potentially important role of the N-methyl-N-benzyl template of 7ACC2 and FACC2, respectively. In this regard, the fluoro-aminocarboxycoumarin derivatives have been designated as lead candidates for further investigation.

### Lead FACC drug candidates specifically inhibit pyruvate driven respiration without substantial effects on other metabolic fuels

Although our initial experiments and previous literature illustrate the ability of the candidate compounds to inhibit pyruvate driven respiration, we sought to investigate the ability of candidate compounds to inhibit respiration fueled by other metabolic substrates glutamate and succinate (**Figure 2**). Glutamate, similar to pyruvate oxidation results in NADH equivalents that fuel complex I mediated respiration, and succinate metabolism offers FADH_2_ for electrons in complex II driven respiratory processes. Hence these metabolites play an important role in the overall viability of oxidative phosphorylation. To further validate the mode of action of the candidate compounds, a series of similar Seahorse experiments were employed (**Figure 2A-C**). In these experiments, 4T1 cells were first permeabilized and were then offered metabolic substrates pyruvate (Pyr/Mal **Figure 2A**), glutamate (Glu/Mal, **Figure 2B**), or succinate (**Figure 2C**) and the resulting OCR was observed. As described previously, malate and DCA were included in the substrate milieu of glutamate respiration experiments. Cells were then offered candidate compounds (1µM) and acute changes in OCR demonstrated that the synthesized compounds specifically inhibited pyruvate driven respiration without substantial effects on glutamate or succinate driven respiratory processes (**Figure 2A-D**). To validate that inhibition of pyruvate respiration is due to mitochondrial pyruvate uptake and not other pyruvate processing enzymes (ie pyruvate dehydrogenase), a parallel experiment using methyl pyruvate (MePyr) was employed (**Figures 2E-F**). MePyr is permeable to the mitochondrial membranes, where in the matrix mitochondrial esterase hydrolysis activity provides pyruvate. Hence, MePyr provides a means to deliver pyruvate to the matrix independent of MPC activity, and reversal of the effects on pyruvate driven respiration in this regard demonstrates specific inhibition of MPC by candidate compounds. These experiments illustrated that pyruvate driven respiration effects of 7ACC2 were completely reversed, indicating that 7ACC2 and other similar fluoro-aminocarboxycoumarins are acute and specific MPC inhibitors (**Figures 2E-F**).

**Figure 2.**
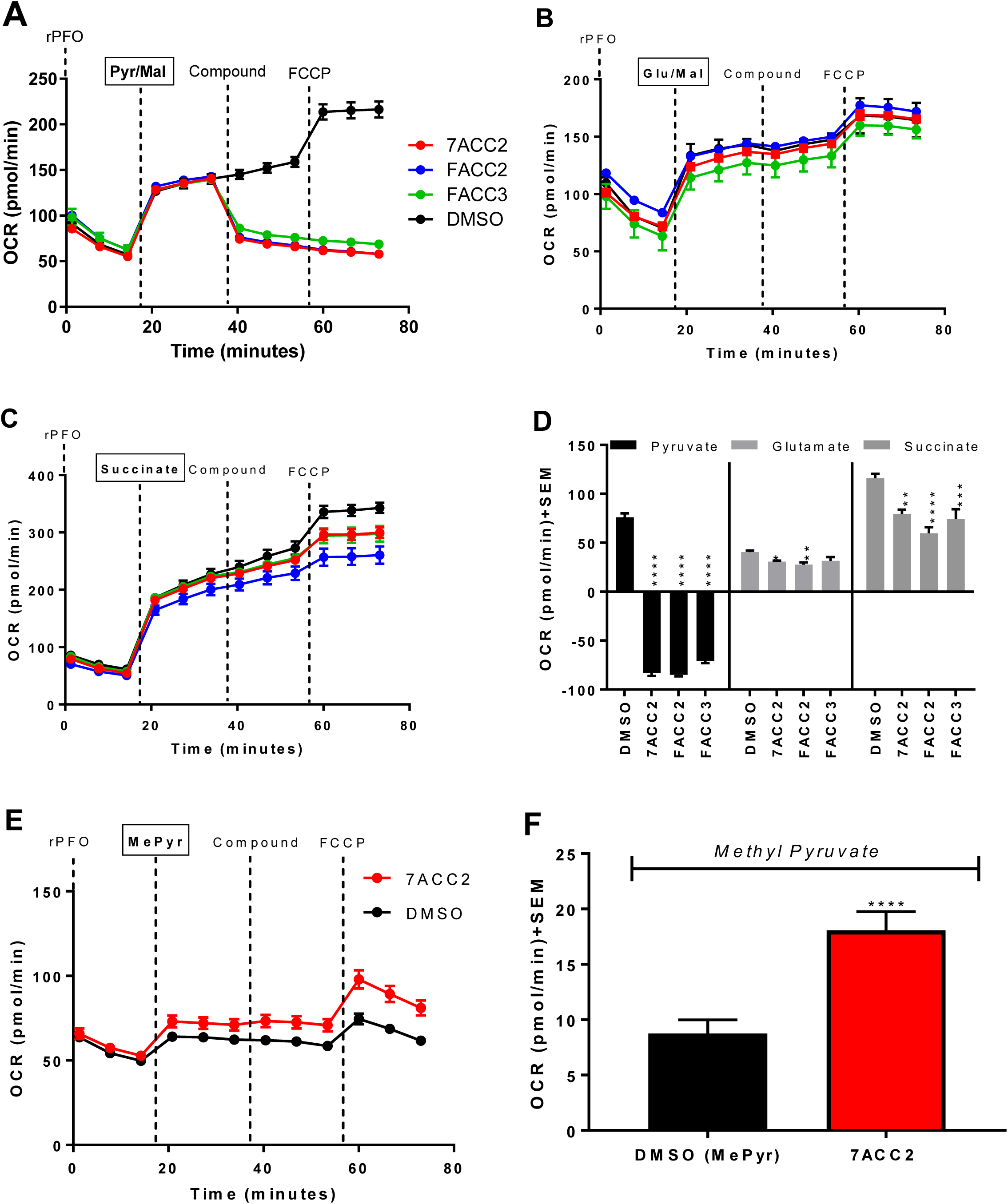
(**A-D**) Candidate compounds specifically inhibit pyruvate driven respiration without substantial effects on glutamate or succinate driven respiratory processes. (**E-F**) Methyl pyruvate reversed the inhibitory capacity of candidate compounds indicating acute MPC inhibition. All experiments are representative of at least three independent experiments, and data represents the average ± SEM. One-way ANOVA analysis was performed to indicate statistical significance between DMSO and compound-treated cultures (*p<0.05, **p<0.01, ***p<0.001, ****p<0.0001).

### Michaelis-Menten kinetics and computational docking studies reveal insights into aminocarboxy coumarin drug candidates MPC binding and inhibitory capacity

To further investigate the type of inhibition, we employed Michaelis-Menton experiments in modified Seahorse XFe96 based assays similar to described above, and based on the model that pyruvate uptake is directly coupled to NADH oxidation and oxygen consumption (**Figure 3A**). In these experiments, 4T1 cells were permeabilized and were then administered varying concentrations of pyruvate in the presence or absence of 10nM 7ACC2 or FACC2, respectively (**Figure 3 B-F**). A rise in OCR indicated pyruvate driven respiration, where subsequent addition of FCCP stimulated maximal respiration and substrate-dependent increases in OCR were observed and were normalized to protein content (**Figure 3 B-F**). In this regard, Michaelis-Menten curves were generated using the FCCP stimulated OCR in 7ACC2 treated and untreated cells where Michealis-Menton parameters of maximal velocity (V_max_) and half-maximal substrate concentrations (K_m_) were calculated (**Figure 3E-G**). Michaelis-Menten assumptions include direct and irreversible substrate and carrier interaction that results in the oxygen consumption rates, and is analogous to standard enzymatic Michaelis Menten models (**Figure 3A**). In the absence of inhibitor, 4T1 cells exhibited a V_max_ of 23.9pmol/min and a K_m_ of 0.07mM whereas in 7ACC2 cells V_max_ and K_m_ values were 10.8pmol/min and 0.19mM respectively (**Figure 3E, F, & H**). Thus, 7ACC2 decreased the Vmax and increased the Km, not-consistent with competitive inhibition where high substrate concentrations titrate out the inhibitor to reach intrinsic Vmax values. However, potential covalent bonding of inhibitor may limit the ability of pyruvate to displace 7ACC2 from the active site, resulting in the observed “mixed” mode of inhibition. Interestingly, FACC2 increased the K_m_ of pyruvate to 0.20mM, similar to 7ACC2, but did not substantially alter Vmax, indicating a similarly potent but unique mode of inhibition (**Figure 3G-H**).

**Figure 3.**
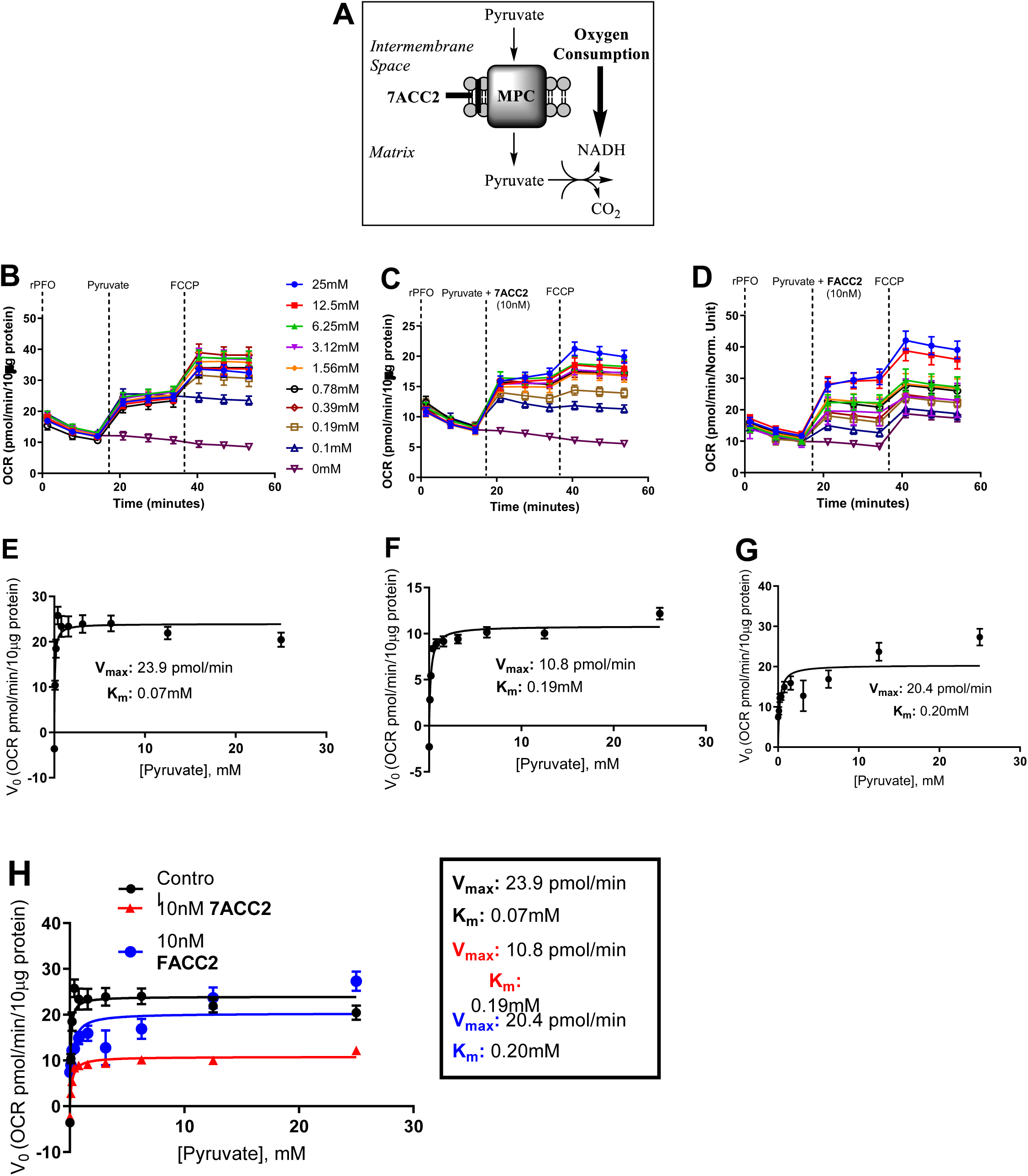
MPC binding characteristics of 7ACC2 are consistent with reversible covalent interactions in with amino acids in the pyruvate binding site. (A-B) Doubly activated olefins of cyanocinamic acid and carboxy coumarin template enable reversible covalent bonding with intracellular nucleophiles. (**C-I**) Seahorse XFe96 assays illustrate michaelis-menten type kinetics of mitochondrial pyruvate uptake, with 7ACC2 demonstrating a “mixed” mode of inhibition based on decreased Vmax and increased Km when compared to control cultures.

To further investigate the inhibitory characteristics of 7ACC2 and FACC2, we carried out computational homology modeling and inhibitor docking studies to give structural insights into binding characteristics on MPC (**Figure 4A-B, Figure S1A-B**). Currently, there are no computationally robust models of MPC available. Here, we generated a model of MPC (open to the cytosol) and subsequent docking studies revealed that the candidate compound interacted with a range of aromatic amino acids and was in close proximity to polar nucleophilic amino acids (**Figure 4B, FigureS1B**). Further, it appeared that ACC2 derivatives bound in regions overlapping with pyruvate binding sites, giving rise to potential competitive modes of action (**Figure 4A**).

**Figure 4.**
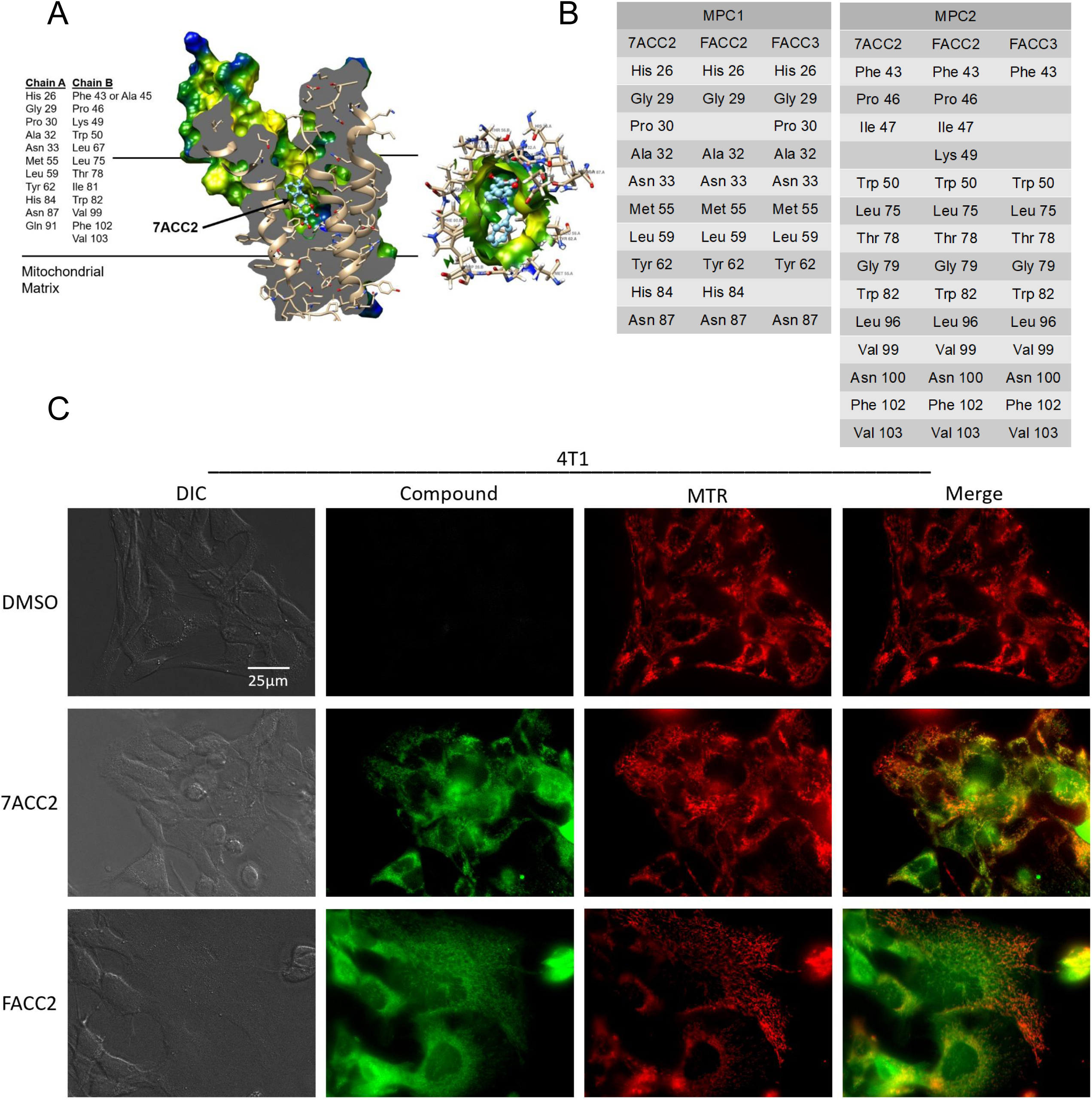
(**A-B**) Computational modeling and inhibitor docking studies of MPC with 7ACC2 reveal intractions with amino acids in the pyruvate binding site. (**C**) Epifluorescent microscopy experiments reveal candidate compounds accumulate in the mitochondria in 4T1 cells. Compound fluorescence was captured under standard GFP filter sets where there was no observed fluorescence overlay into the red (MTR) channel. Images are representative of overall culture appearances (>5 fields of view) and of three independent experiments. All images were captured at the same magnification (see scale bar).

### Epifluorescent microscopy experiments indicate 7ACC2 and FACC2 co-localize with the mitochondria

To further investigate cellular localization of candidate compounds, epifluorescent microscopy experiments were employed (**Figure 4C**). Conveniently, the aminocarboxycoumarin template offers fluorescent capabilities when excited in standard GFP excitation and emission settings and hence, the cellular localization of 7ACC2 and FACC2 candidates can be observed. In this regard, 4T1 cells were seeded in MatTek glass-bottom dishes and were incubated for 48 hours for adherence and flattening. Cells were then exposed to either 7ACC2 or FACC2 for 30min, followed by Mitotracker-RED CMXROS (MTR) for 15 min at 37°C. Growth media was then aspirated and replaced with MAS buffer + 5% fetal bovine serum for imaging. Here, we observed that candidate compounds accumulated in granular structures, and significant overlay with MTR indicated substantial accumulation in the mitochondria consistent with MPC inhibition (**Figure 4C**).

### Water soluble and metabolically stable aminocarboxycoumarin MPC inhibitor D7 exhibits enhanced water solubility and metabolic stability and elicits anticancer properties in models of breast cancer

To improve the drug like properties of lead candidate FACC2, we sought to synthetically modify 1) the fluoro substitution with more lipophilic and metabolically stable trifluoromethyl group (compound **11**, **Scheme 2A**) and 2) introduce a water-soluble carboxylate salt by synthesizing **D7** (**Scheme 2B**). The synthesis of **D7** began with silylation of 3-aminophenol 1 with ^t^butyldiphenylsilyl chloride **2** in the presence of imidazole base in DCM (**Scheme 2A**). This newly protected amine **3** was then subjected to reductive amination by sodium borohydride after condensation with 4-(trifluoromethyl)benzaldehyde **4** in refluxing ethanol. The aryl mono substituted silyl protected amine **5** was then methylated using methyl iodide in DMF to yield **6**. This disubstituted amine **6** was then deprotected in THF using HCl. The newly formed phenol **7** was subjected to Vilsmeier-Haack formylation using POCl3 in DMF. This salicylaldehyde **8** then underwent Knoevenagel condensation with diethyl malonate **9** in the presence of piperidine in ethanol. The condensation product **10** then underwent NaOH hydrolysis followed by acidification and cyclization using HCl to afford **11**, the carboxylic acid form of **D7** in ∼72% yield (**Scheme 2A**).

One limitation of N,N-dialkylaminocoumarin derivatives is a lack of water solubility. To mitigate this issue, we sought to synthesize a salt of **D7** with high solvating capacity (**Scheme 2B**). Starting with the carboxylic acid form **11** of **D7**, various salt forms were generated to increase the solubility **D7**. A sodium salt was first generated but did not substantially increase the water solubility. Incorporating organic salt forms in drug development offers additional hydrogen bonding acceptor or donor groups and significantly enhances the solubility of organic compounds. Tris(hydroxymethyl)aminomethane (tris base, **12**) was utilized in the further chemical optimization of **D7**. Tris base **12** was utilized due to its high solubility in water, low toxicity, and due to its pKa being slightly above physiological pH (pKa = 8.07), providing the ammonium counter-ion at physiological pH. The carboxylic acid form 11 of D7 was dissolved in methanol followed by the addition of tris base **12**. After stirring for 1 hour a bright yellow solid precipitated out, filtered, and rinsed with cold MeOH to yield the tris base form, D7 (**Scheme 2B**).

To confirm that D7 retained mitochondrial targeting capacity, we first carried out standard Seahorse mitochondrial stress tests, where assay media contains a variety of metabolic fuels including glucose, pyruvate, and glutamine (**Figure 5A-E**). Here, we observed that D7 inhibited mitochondrial respiration in a dose-dependent fashion with notable decreases in oxygen consumption upon acute injection of compound (**Figure 5C**), following oligomycin injection indicating decreases in ATP production (**Figure 5D**), and following FCCP illustrating decreases in maximal respiration (**Figure 5E**). Interestingly, we observed a simultaneous and dose-dependent decrease in compensatory glycolysis (extracellular acidification rates, ECAR), suggesting a potential feedback inhibition of glycolysis in the MCT1 expressing 4T1 cells, consistent with MPC inhibition. Thus, we further characterized D7 toward the inhibition of pyruvate driven respiration as described in **Figures 1** & **2**. Here, we found that D7 inhibits pyruvate-driven respiration to a similar extent when compared to both 7ACC2 and FACC2 with and IC50 value of 14.4 ± 0.5 nM in permeabilized 4T1 cells (**Figure 5F-G**). Consistent with FACC2, **D7** did not alter mitochondrial respiration when offered glutamate or succinate (**Figure S2A-D**). Further, methyl pyruvate was able to reverse the D7-inhibited pyruvate driven respiration, indicating that this inhibition is transport mediated (**Figure S2E-H**). Taken together, **D7** perturbs mitochondrial respiration consistent with MPC inhibition, qualifying this drug candidate for further anticancer efficacy studies.

**Figure 5.**
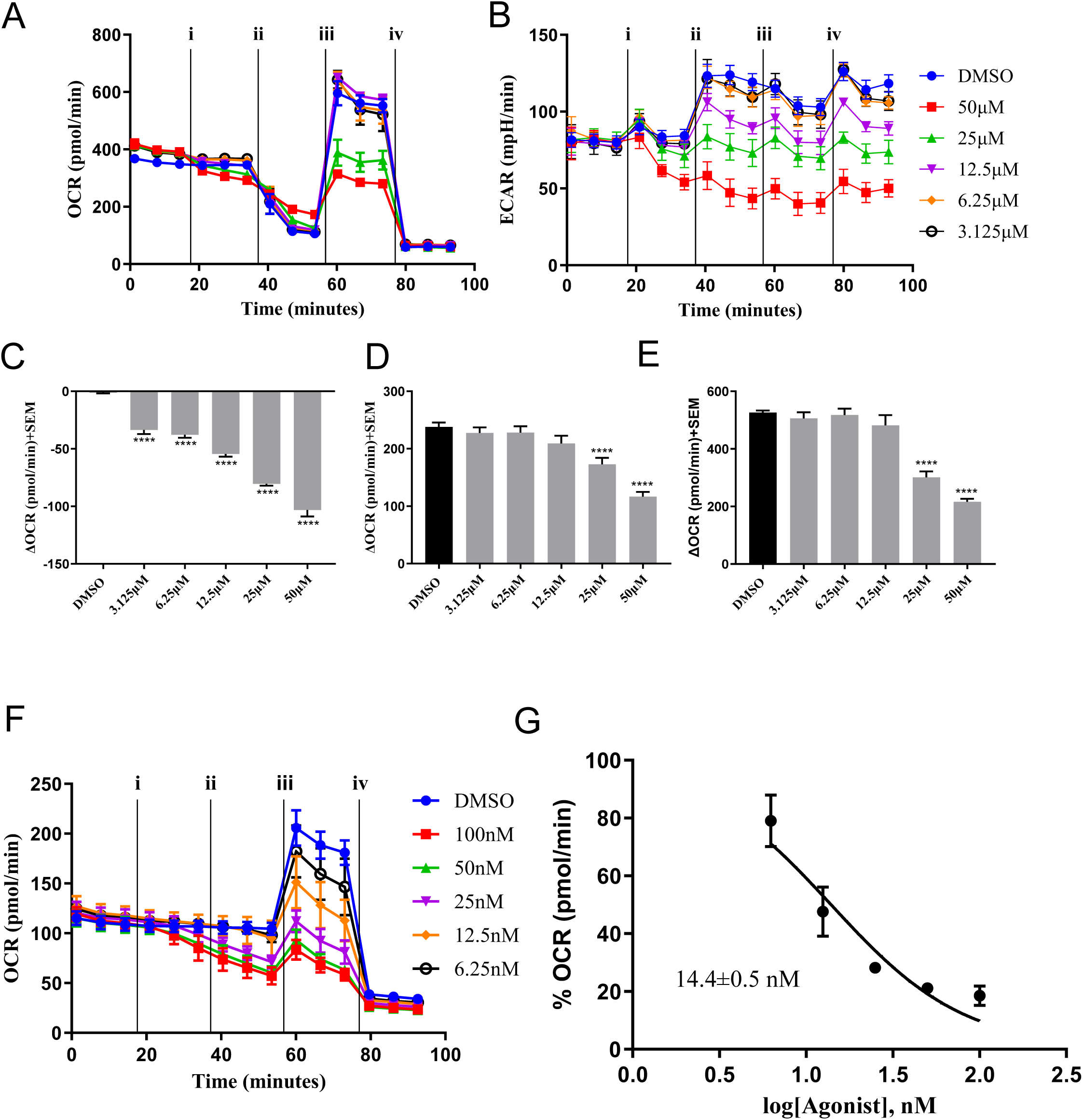
Seahorse-based mitochondrial stress test illustrates that D7 inhibits (**A**) mitochondrial respiration (oxygen consumption rates, OCR) and (**B**) compensatory glycolysis (extracellular acidification rates, ECAR) in in-tact 4T1 cells. OCR and ECAR were measured following injection of (i) D7, (ii) oligomycin, (iii), FCCP, and (iv) a cocktail or rotenone and antimycin A. D7 reduces (**C**) acute OCR, (**D**) ATP-production, and (**E**) maximal respiration. (**F-G**) D7 inhibits pyruvate driven respiration in permeabilized 4T1 cells.

As noted previously, D7 treatment resulted in a dose dependent decrease in glycolysis which may be due to cytosolic accumulation of lactate upon MPC inhibition (**Figure 6A**). To test this, we assayed intracellular lactate accumulation following D7 treatment in MCT1 expressing 4T1 cells, and MCT4 expressing MDA-MB-231 cells using commercially available lactate kits. MCT1 mediates lactate import, and MCT4 mediates lactate export, where 4T1 and MDA-MB-231 cells offer two cellular models of lactate accumulation, where MCT4 expression is predicted to alleviate lactate accumulation upon MPC inhibition. Our results indicate that D7 treatment results in a potent accumulation of lactate in 4T1 (**Figure 6B**), but not MDA-MB-231 (**Figure 6C**) cells, consistent with our hypothesis. It is possible that an intracellular accumulation of lactate can contribute to cell proliferation inhibition properties of D7 in addition to its mitochondrial targeting capacity. Interestingly, consistent with lactate accumulation experiments, D7 is selectively toxic to 4T1 and isogeneic 67NR cell lines when compared to MDA-MB-231 (**Figure 6D**), indicating that MPC inhibition may be particularly effective in MCT1 expressing contexts.

**Figure 6.**
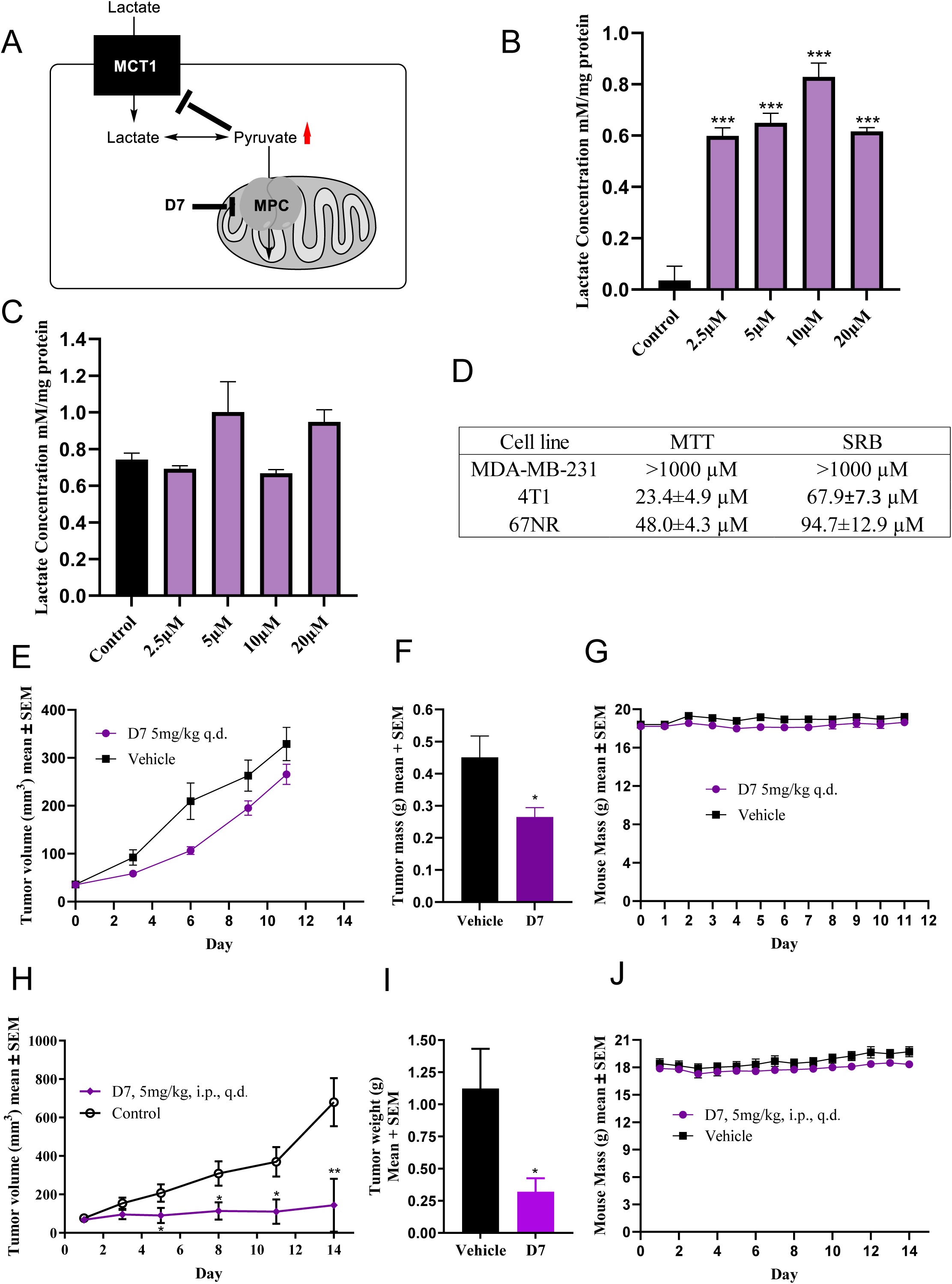
D7 alters perturbs lactate flux and exhibits anticancer efficacy in *in vitro* models of breast cancer. (**A**) Model illustrating that inhibition of pyruvate uptake into mitochondria results in accumulation of intracellular lactate. D7 treatment results in the accumulation of lactate in (**B**) 4T1 but not (**C**) MDA-MB-231 cells. (**D**) D7 is specifically toxic in 4T1 and 67NR when compared to MDA-MB-231. (E) D7 inhibits 4T1 tumor volume and (F) tumor mass without affecting (G) mouse body weight. (H-J) D7 exhibits anticancer efficacy in a 67NR tumor model.

To further evaluate the anticancer efficacy of D7, we performed *in vivo* syngraft models using two isogeneic murine cell lines 4T1 and 67NR derived from distinct sites from the same mouse. 4T1 cell line is characterized as highly metabolically plastic and aggressive, whereas 67NR is less-so. For this experiment, 0.1 mL of 4T1 cells were injected into the flanks of BALB/c mice with a 1:1 mixture of Matrigel and PBS containing 12,500 cells/mouse. After tumors reached 50 mm^3^, mice were randomly placed into groups. **D7** was then administered as a 10% DMSO/water solution at 5 mg/kg daily for 11 days. After 11 days, vehicle control mice began to ulcerate so mice were sacrificed, and tumors were resected. After 11 days of treatment, it was found that **D7** treated mice saw a ∼20% reduction in tumor volume compared to vehicle treated control (**Figure 6E**). After tumor resection it was found that D7 treated groups caused a ∼41% reduction by mass when compared to vehicle treated control (**Figure 6F**). Mice weights remained constant throughout the duration of the study (**Figure 6G**). For the 67NR model, 500,000 67NR cells in a 1:1 solution of matrigel and PBS for a total volume of 0.1 mL were administered into the flank of BALB/c mice. Tumors were allowed to grow until they were ∼100mm^3^ and randomly grouped. At that time, **D7** was administered 5 mg/kg intraperitoneally once daily for 14 days. After 14 days, there was a ∼79% reduction in tumor volume in D7 groups as compared to vehicle treated (**Figure 6H**). At the end of the study, tumors were resected, and tumor mass was recorded. Compared to vehicle control, D7 was found to reduce tumor mass by ∼71% (**Figure 6I**). Mouse body weights remained stable during the 14-day treatment period (**Figure 6J**). Gratifyingly, MPC inhibition characteristics where conserved in the 67NR cell line (**Figure S3A-B**).

Pharmacokinetic parameters of **D7** in BALB/c mice were evaluated using three mice per time point, plasma samples were collected after IV treatment of 1 mg/kg and PO treatment of 5 mg/kg of **D7** in both male and female mice (**Figure S3C**). Using this data, pharmacokinetic parameters such as half-life (t_1/2_), T_max_, C_max_, apparent volume distribution at steady state (V_ss_), clearance (CL), and bioavailability using Pheonix WinNolin 7.0 in a noncompartmental analysis method. IV dosing allows for analysis under the assumption of total systemic absorption (**Figure S3C**). Under this assumption it was found that D7 has a t_1/2_ of 3.15 and 3.65 hours with CL of 26.61 mL/min/kg and 34.30 mL/min/kg in female and male mice respectively. D7 was found to have V_ss_ of 6.92 L/kg in female mice and 10.24 L/kg in male mice. Volumes of distribution above 5 L/kg are considered medically to be high values, indicating that D7 has a high V_ss_ in both male and female mice and may have a likelihood to leave plasma and distribute into tissues.177 However, with a noncompartmental analysis method, the tissue specific accumulation of D7 is not known at this time. In oral dosing of D7, the t_1/2_ was found to be 4.41 and 4.04 hours in female and male mice respectively. Additionally, the bioavailability was found to be 124.7% and 123.8% in female and male mice respectfully (**Figure S3C**). It is not possible to obtain a bioavailability over 100%, however, we can extrapolate these values to mean that D7 has close to 100% bioavailability. These studies illustrate that D7 exhibits tumor growth inhibition properties in a variety of tumor models, and exhibits favorable pharmaceutical properties including water solubility and metabolic stability to be considered as a suitable anticancer agent for further pre-clinical development.

## CONCLUSIONS

In conclusion, we have synthesized and evaluated a novel series of fluoro-substituted *N,N-* dialkylcyanocinnamic acid and aminocarboxycoumarin derivatives with potent MPC inhibition properties. Our studies indicated that the aminocarboxycoumarin template was superior to *N,N-* dialkylcyanocinnamic acid in eliciting MPC inihibitory characteristics, and specifically, structure activity relationship studies revealed that the *N-*methyl-*N-*benzyl structural template offered the optimal inhibitory capacity. Further respiratory experiments illustrated that candidate compounds specifically inhibit pyruvate driven respiration without substantially affecting other metabolic fuels consistent with MPC inhibition. Further, aminocarboxycoumarin binding characteristics were found to be consistent with reversible covalent bonds with aminoacids in the pyruvate binding domain. Epifluorescent microscopy experiments illustrated that FACC2 accumulates in the mitochondria to a similar extent as parent 7ACC2. We have further developed these first generation inhibitors by improving upon water solubility and pharmaceutics by synthesizing **D7** which exhibits anticancer efficacy in a variety of tumor models.

## MATERIALS AND METHODS

### Synthesis of fluoro-aminocarboxycoumarin derivatives and D7

Synthetic procedures and compounds characterization are included in the supplementary information.

### Seahorse XFe96 based respiratory experiments

Permeabilized cell assays were performed using rPFO as described previously. 4T1 cells were seeded (20,000cells/well) onto Seahorse XFe96 well plates and incubated overnight in growth media at 37C and 5%CO_2_ for adherence. On the day of the assay, growth media was aspirated and replaced with mannitol/sucrose buffer (MAS; 70mM sucrose, 220mM mannitol, 10mM potassium phosphate monobasic, 5mM magnesium chloride, 2mM HEPES, and 1mM EGTA) after 3X rinse of growth media to remove serum and endogenous metabolic substrates, and incubated at 37C in a non-CO_2_ incubator. Respective inhibitor and substrate milieu’s were prepared in MAS buffer for port injections A-D at 8X, 9X, 10X, and 11X the target cell concentrations to account for intrinsic dilution factor of *in situ* injections of each port. For some experiments (Figure 4A&I), compound **2** was injected in port A, followed by rPFO (1nM) in port B, followed by respective substrate cocktails (FCCP stimulated) in port C, and rotenone and antimycin A (0.5µM) in port D. In other experiments (Figure 4C-H), permeabilization was initiated prior to port A injection during the MAS buffer wash phase, followed by substrates, test compound, and FCCP (0.125 µM). Final substrate concentrations for specific tests were as follows: (5mM pyruvate, 0.5mM malate, 2mM dichloroacetate (DCA); 10mM glutamate, 0.5mM malate, 2mM DCA; 10mM succinate, 2µM rotenone; 20mM methyl pyruvate, 5mM pyruvate, 0.5mM malate, and 2mM DCA). ATP rate assays and mitochondrial stress tests were performed according to the manufactures (Agilent) instructions.

### Western blotting

4T1 cells were seeded in 100mm dishes (5×10^5^ cells/dish) and were incubated for 24 hours and whole cell lysates were collected in RIPA buffer, and large proteins were precipitated and discarded via centrifugation. After denaturation at 95°C for 10 min in RIPA buffer, protein concentration was determined and equilibrated using the BCA assay (ThermoScientific) and 20 μg of protein was separated by electrophoresis in 12% polyacrylamide gels. Proteins were electrophoretically transferred to nitrocellulose membranes for 60 min at 100 V and stained with Ponceau to verify transfer quality and equal protein loading. After blocking the membranes with 10% w/v nonfat milk in PBST for 1 h at 35⁰C, membranes were incubated overnight at 4°C with primary antibodies (MPC1, abcam, ab74871; MPC2, abcam, ab236584, 1:500; PBST, 1% nonfat milk protein). Membranes were rinsed three times with PBST and incubated with the respective secondary goat anti-mouse IgG (1:2000) or goat anti-rabbit IgG (1:2000) horseradish peroxidase-conjugated antibodies (Jackson Immunoresearch). Membranes were again washed three times with PBST and exposed to SuperSignal® 60 West Pico luminol enhancer and stable peroxide solution (ThermoScientific) for 2 minutes. Chemiluminsecent images were obtained using a LicorFC imager. Images shown are representative of three separate experiments.

### Computation homology modeling and inhibitor docking studies

Structures were generated for human MPC1 and 2 by homology modeling with MODELLER 9.18. Autodock Vina was used to dock 7ACC2 to the outward (cytosol) open homology models. Experimental protocols were achieved in a similar fashion as described by us previously.

### Epifluorescent microscopy

4T1 cells (5×10^4^ cells/mL) were seeded in glass-bottom dishes (MatTek Corp, part no. P35G010C) and incubated for 48 hours. 7ACC2 or FAAC2 were then added (10μM) and cells were again incubated for 6hr 30 minutes and MitoTracker Red CMXROS (Invitrogen, M7512, 100nM) was added 15min prior to microscopic fluorescent imaging. Media was then aspirated and replaced with MAS+5% FBS for imaging. Cells were then examined and photographed using a Nikon TE2000 epifluorescent microscope and camera. The images shown are representative of at least 5 fields of view of three separate experiments and captured under the same 60X oil emersion DIC lens, see scale bar.

### L-lactate accumulation assay

MDA-MB-231 or 4T1 cells were seeded on a 6-well plate at 150,000 cells/well in their respective culture medias and incubated for 3 days (37°C, 5% CO_2_). On the third day, media was aspirated, and wells were rinsed twice with cold PBS. After rinsing, phenol red free RPMI-1640 (10% FBS and penicillin-streptomycin (50 μg/mL)) was added with five treatment concentrations and one control and incubated for 24 hours (37°C, 5% CO_2_). After 24 hours, extracellular media was removed, and cells were washed 3 times with ice cold PBS. At this time, 1 mL of PBS was added to each well and underwent freeze-thaw lysis at -80°C. Media in each well was aliquoted and centrifuged (16,000 x g, 10 minutes, 4°C). The lysate supernatant was then aliquoted, and pellet discarded. L-lactate concentration was then determined using Eton Biosciences L-Lactate assay kit 1 by mixing lysate sample with L-lactate assay solution in duplicate, incubating for 30 minutes (37°C, 0% CO_2_) and measuring absorbance of enzyme produced INT formazan at 490 nm using a BioTek Synergy 2 platereader. Absorbance values were compared to a L-lactate standard curve concomitantly ran with test samples to determine lactate concentration in each sample. Lactate concentrations were normalized to total protein using a Pierce BCA Protein Assay Kit.

### MTT Cell Proliferation Inhibition Assay

Confluent cell cultures were treated with trypsin and resuspended at 5×10^4^ cells/mL. To a 96-well plate, 100 µl of the 5×104 cells/mL solution were added and allowed to incubate at 37°C, 5% CO_2_ for 24 hours. Compounds were then added and allowed to incubate for 3 days. At this time 10 µL of MTT (5 mg/mL) was added to the 96-well plate and further incubated for 4 hours. Following the 4-hour incubation with MTT, 100 µL of SDS (0.1 g/mL, 0.01N HCl) was added and 96-well plates were allowed to incubate for an additional 4 hours. Absorbance values were then taken at 570 nm using a Biotek Synergy 2 plate reader. Treatment absorbances are taken as percentages of the average untreated absorbances and plotted against log(concentration) in GraphPad Prism 10 to generate effective concentration values were 50% of the cells are not proliferating (EC_50_).

### SRB Cell Proliferation Inhibition Assay

Cells were seeded (3×10^4^cells/well) in 48 well plates and incubated overnight (37°C, 5%CO_2_). Test compounds were then exposed to cells in duplicate in serial dilution fashion and were incubated for an additional 72 hours (37°C, 5%CO_2_). Media was then aspirated, and cells were washed three times with cold PBS and left overnight to dry fix. 100µL Sulforhodamine B (SRB, 0.5% w/v in 1% aqueous acetic acid) was added to each well and was incubated at 37°C for 1hr. SRB was rinsed with 1% acetic acid solution and dried. Dyed cells were then lysed with Tris (10mM, pH 10) and absorbance readings were recorded at 540nm of each well. Treatment absorbances are taken as percentages of the average untreated absorbances and plotted against log(concentration) in GraphPad Prism 10 to generate effective concentration values were 50% of the cells are not proliferating (EC50).

### Tumor efficacy study in a 67NR syngraft

Female BALB/c (Charles River) were administered 67NR at 500,000 cells/mouse into their flank as a 1:1 solution of Matrigel and PBS for a total volume of 0.1 mL. Tumors were grown until they reached a volume of ∼100 mm^3^ and were then randomized into groups (n=6). Mice were administered intraperitoneally either vehicle control (10% DMSO, 90% sterile water) or D7 at 5 mg/kg once daily for 14 days. Body weights were recorded daily and monitored for behavior and grooming patterns. A body weight loss (BWL) >10% indicated a halting of treatment until weight recovery. If a BWL >20% was observed, that mouse would be euthanized. Tumor volume (TV, mm^3^) was calculated as follows: TV = (a×b^2^)/2, a as the tumor length and b as the tumor width. The tumors were measured every 2-3 days using a caliper in two dimensions. Mice were euthanized at the end of the 14-day treatment period where tumors were then resected and weighed. The study was carried out under IACUC protocol 2106-39167A. Statistical analysis was carried out via unpaired t-test using GraphPad Prism 10 (*p<0.05, **p<0.01, ***p<0.001, ****p<0.001).

### Tumor efficacy study in a 4T1 syngraft

Female BALB/c (Charles River) were administered 67NR at 12,500 cells/mouse into their flank as a 1:1 solution of Matrigel and PBS for a total volume of 0.1 mL. Tumors were grown until they reached a volume of ∼50 mm^3^ and were then randomized into groups (n=6). Mice were administered intraperitoneally either vehicle control (10% DMSO, 90% sterile water) or D7 at 5 mg/kg once daily for 11 days. Body weights were recorded daily and monitored for behavior and grooming patterns. A body weight loss (BWL) >10% indicated a halting of treatment until weight recovery. If a BWL >20% was observed, that mouse would be euthanized. Tumor volume (TV, mm^3^) was calculated as follows: TV = (a×b^2^)/2, a as the tumor length and b as the tumor width. The tumors were measured every 2-3 days using a caliper in two dimensions. Mice were euthanized at the end of the 14-day treatment period where tumors were then resected and weighed. The study was carried out under IACUC protocol 2106-39167A. Statistical analysis was carried out via Dunnett’s one-way ANOVA using GraphPad Prism 10 (*p<0.05, **p<0.01, ***p<0.001, ****p<0.001).

### Pharmacokinetic assessment of D7 in a noncompartmental analysis model using BALB/c mice

In a contract organization Medicilon (Beijing, China), 35 male and 35 female BALB/c mice were randomized and put into 2 groups (n=15) per sex for each route of administration (4 total groups). Extra mice were used for collection of blank plasma. Ten designated time points were utilized of 0.083, 0.25, 0.5, 1, 2, 4, 6, 8, 10, and 24 hours. D7 (10% DMSO, 90% sterile water) was administered IV at 1 mg/kg (5 mL/kg) via tail vein injection (n=3 per arm) and orally via gavage after overnight fasting at 5 mg/kg (10mL/kg) (n=3 per arm). After administration of D7, 100 µL of blood was taken from the submandibular or other suitable vein at the designated time points, cardiac puncture was used for terminal bleeding. Each time point utilized three mice per group and each mouse had two blood draws. The blood samples were collected in K_2_EDTA tubes on ice, centrifuged at 6,800 x g (6 mins, 4°C) to generate plasma samples. The plasma samples were stored below -70°C until LC-MS/MS analysis. The LC-MS/MS method development and sample analysis were performed by Medicilon testing facility and the accuracy of >66.7% of the quality control samples should be within 80-120% of their nominal values. After data acquisition, noncompartmental analysis parameters were calculated by Pheonix WinNonlin 7.0.

### Ethical considerations

The general tolerability study in CD-1 was approved and in compliance with the University of Minnesota’s Institutional Animal Care and Use Committee [2211-40546A]. This study was performed in accordance with the relevant guidelines and regulations, and all protocols were approved by the University of Minnesota. The BALB/c general tolerability study with oral dosing was carried out by a contract organization Medicilon Incorporated. Animals were purchased from Beijing Vital River Laboratory Animal Technology Co., Ltd. (Beijing, China). The study was carried out using license No.: SCXK (Beijing) 2021-0006 and animal Certificate No.: 110011220106449863. The 67NR and 4T1 tumor syngrafts were approved and in compliance with the University of Minnesota’s Institutional Animal Care and Use Committee [2106-39167A]. This study was performed in accordance with the relevant guidelines and regulations, and all protocols were approved by the University of Minnesota.

## Supplementary Figures

**Figure S1A-B.**
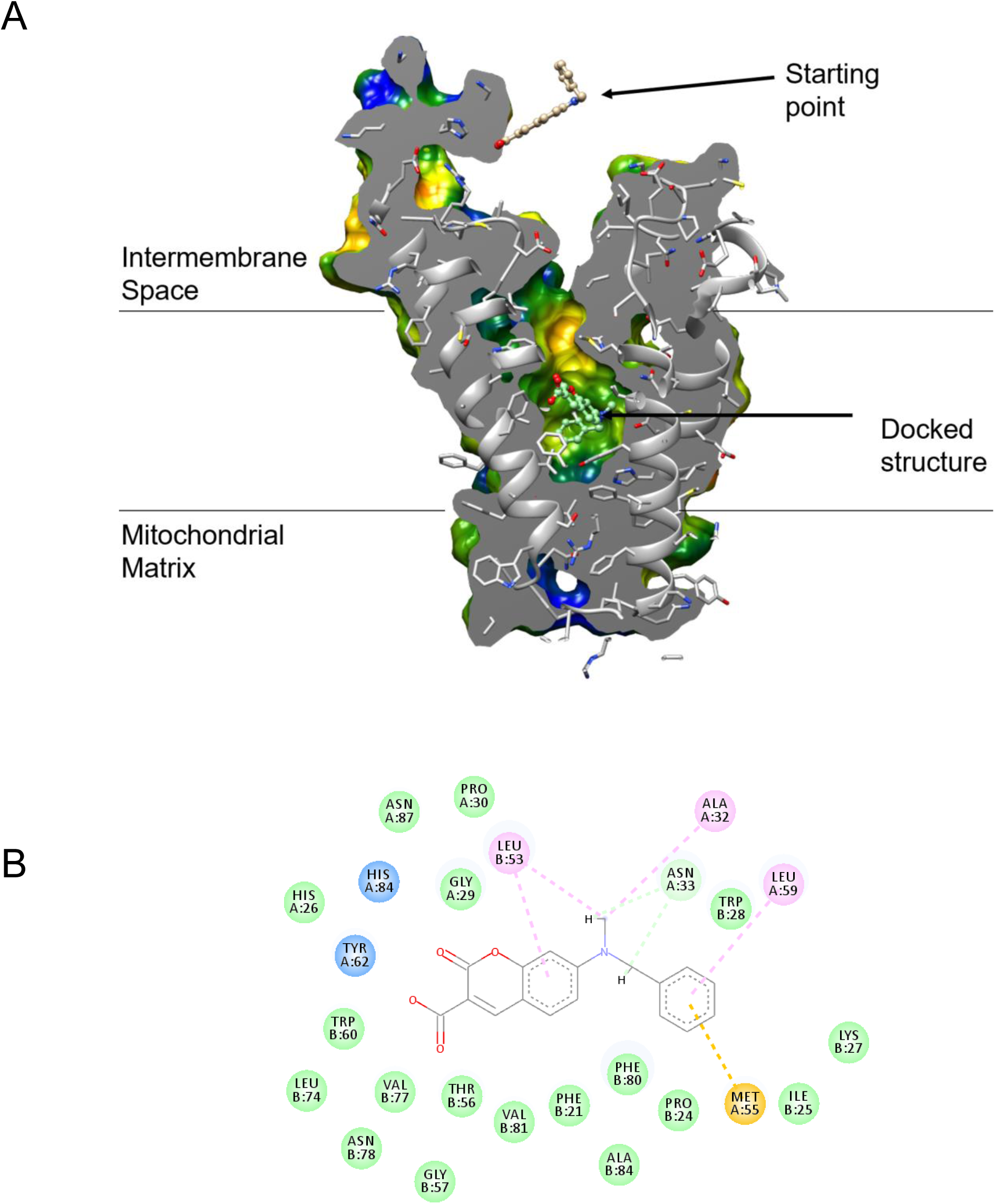
Computational modeling and inhibitor docking studies.

**Figure S2.**
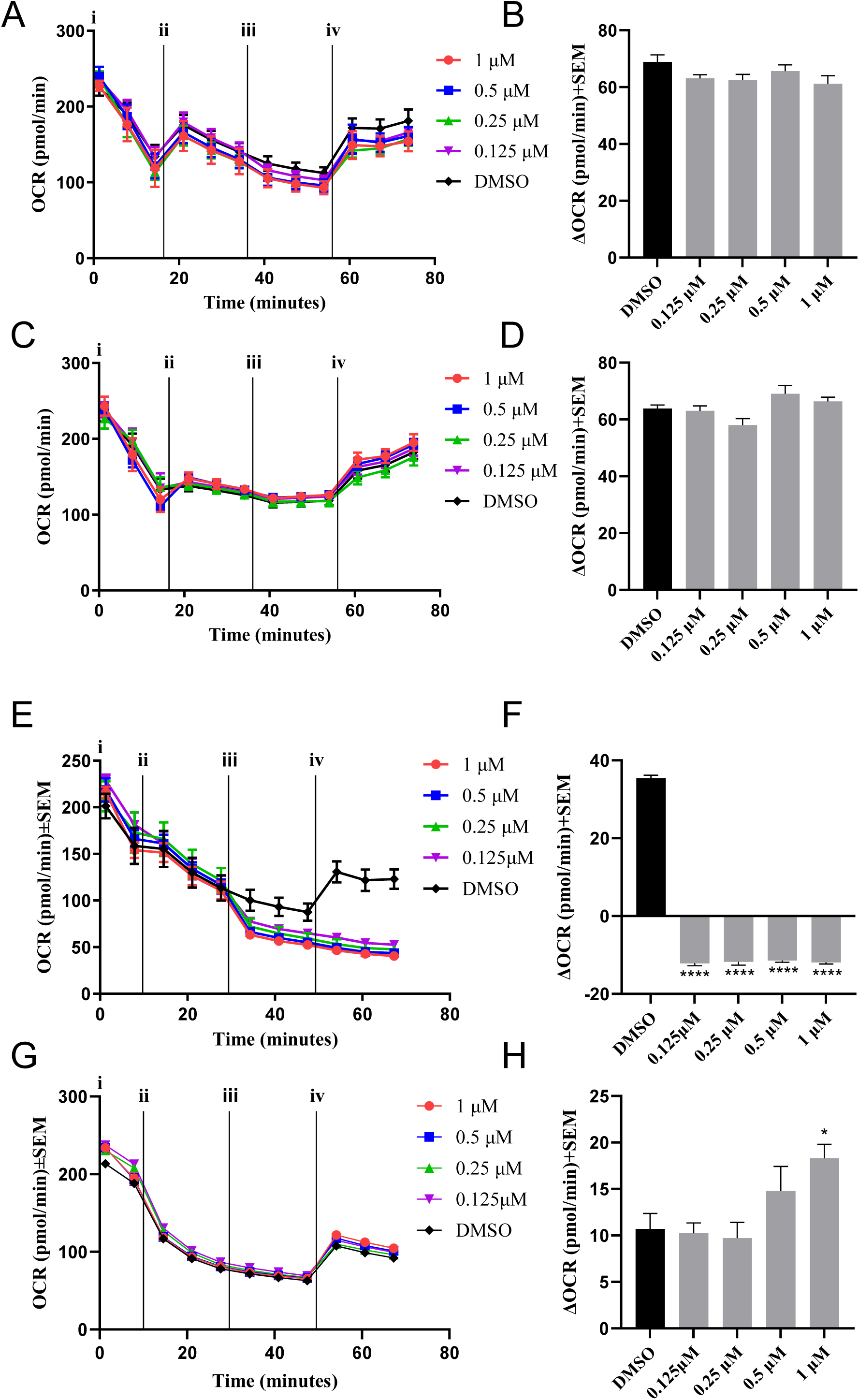
(**A-B**) D7 does not alter glutamate driven respiration. (**C-D**) D7 does not alter succinate driven respiration. (**E-H**) Methyl pyruvate reversers D7-inhbited pyruvate driven respiration.

**Figure S3.**
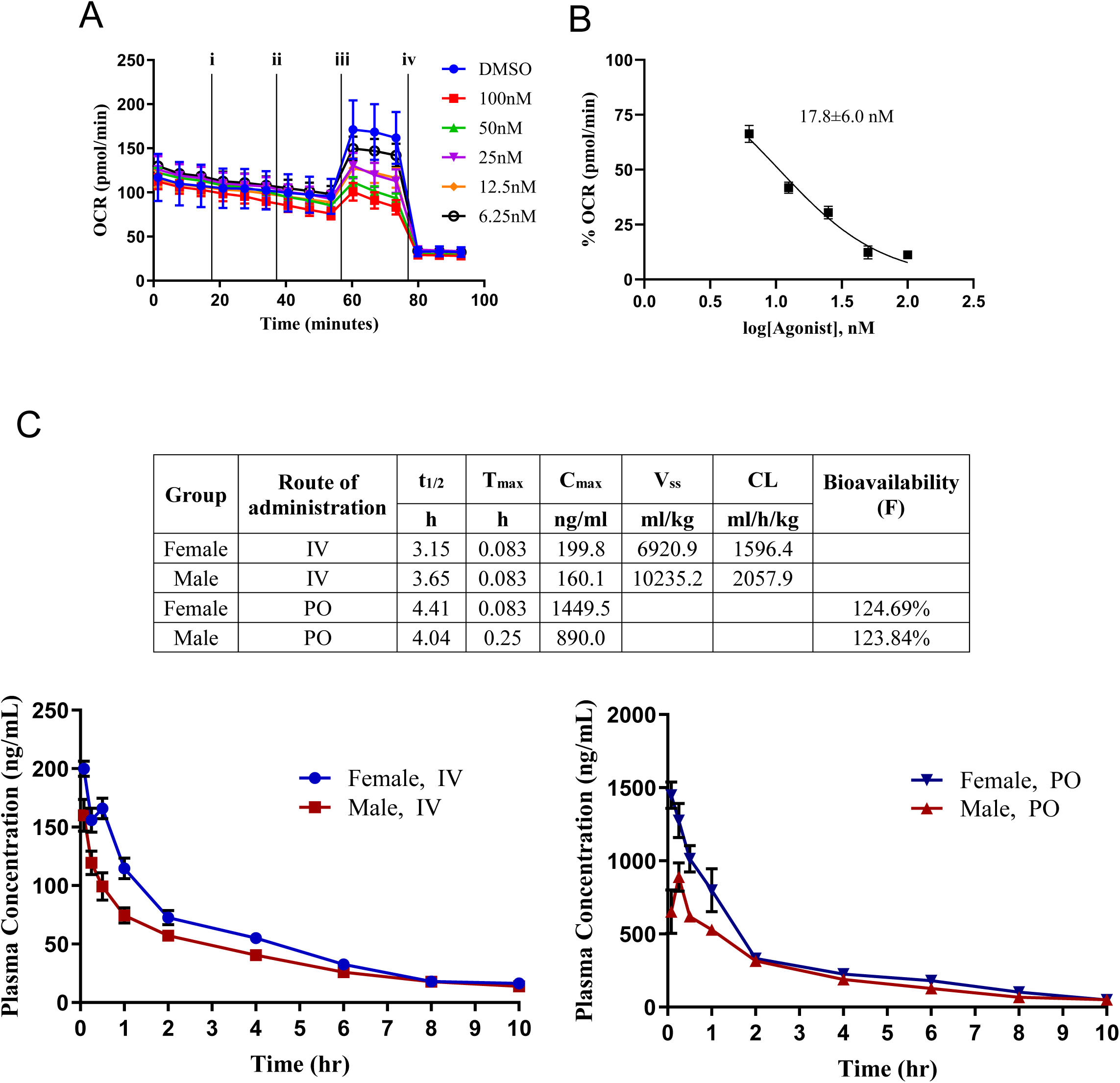
(**A-B**) D7 inhibits pyruvate driven respiration in permeabilized 67NR cells. (**C**) Pharmacokinetic properties of D7 in male and female mice.

## APPENDIX

### Spectral Characterization

^1^H- and ^13^C-NMR spectra were plotted on a Bruker-400MHz NMR or Varian-500MHz NMR. High-resolution mass spectra (HRMS) were recorded using a Bruker BioTOF II ESI mass spectrometer.

*7-(bis(4-fluorophenyl)amino)-2-oxo-2H-chromene-3-carboxylic acid **(FACC1)***

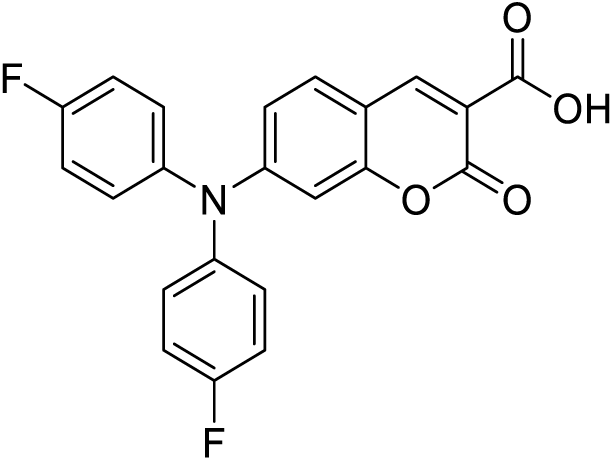

**1H NMR (400 MHz, CHLOROFORM-d):** δ 8.72 (s, 1H), 7.45 (d, J= 8.84 Hz, 1H), 7.23-7.20 (m, 4H), 7.15-7.11 (m, 4H), 6.83 (dd, J= 2.28, 8.84 Hz, 1H), 6.70 (d, 2.16 Hz, 1H)

**13C NMR (100 MHz, CHLOROFORM-d):** δ 164.81, 163.63, 161.02 (d, J_CF_ = 246.9 Hz), 156.95, 154.84, 150.44, 140.46 (d, J_CF_ = 3.05 Hz), 131.45, 128.68 (d, J_CF_ = 8.49 Hz), 117.32 (d, J_CF_ = 22.9 Hz), 115.84, 111.42, 109.02, 103.42

**HRMS (ESI) m/z:** calc’d for C_22_H_13_F_2_NO_4_ [M+H^+^]: 394.0885 found: 394.0917

*7-((4-fluorobenzyl)(methyl)amino)-2-oxo-2H-chromene-3-carboxylic acid **(FACC2)***

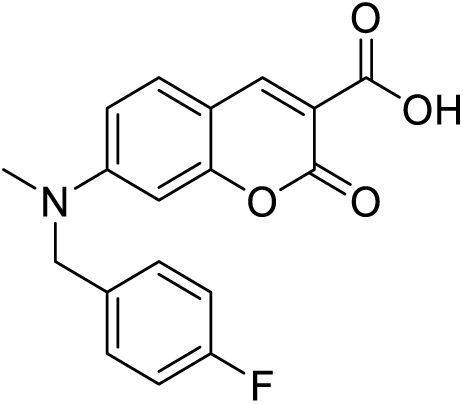

**1H NMR (400 MHz, CHLOROFORM-d):** δ 12.28 (s, 1H), 8.65 (s, 1H), 7.46 (d, J= 9.00 Hz, 1H), 7.16-7.13 (m, 2H), 7.05 (t, J= 8.52 Hz, 2H), 6.75 (dd, J= 2.28, 8.96 Hz, 1H), 6.58 (d, J= 2.08Hz, 1H), 4.68 (s, 2H), 3.23 (s, 3H).

**13C NMR (100 MHz, CHLOROFORM-d):** δ 165.29, 164.09, 162.31 (d, J_CF_ = 245.19 Hz), 157.64, 155.15, 150.50, 131.88, 131.54 (d, J_CF_ = 3.23 Hz), 127.92 (d, J_CF_ = 8.09 Hz), 116.09 (d, J_CF_ = 21.55 Hz), 111.28, 109.21, 106.85, 97.77, 55.66, 39.31.

**HRMS (ESI) m/z:** calc’d for C_18_H_14_FNO_4_ [M+H^+^]: 328.0980 found: 328.0994

*7-((4-fluorophenyl)(methyl)amino)-2-oxo-2H-chromene-3-carboxylic acid **(FACC3)***

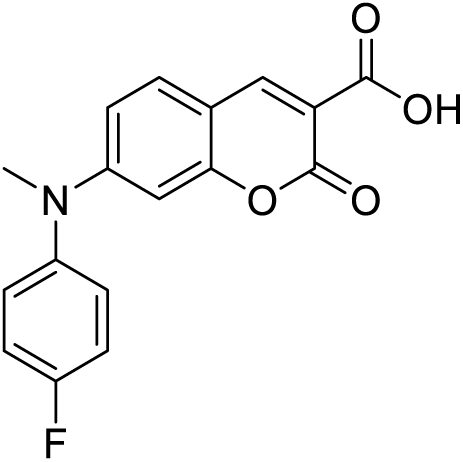

**1H NMR (400 MHz, CHLOROFORM-d):** δ 12.24 (s, 1H), 8.66 (s, 1H), 7.41 (d, J= 8.92 Hz, 1H), 7.27-7.17 (m, 4H), 6.65 (dd, J= 1.4, 8.9 Hz, 1H), 6.57 (s, 1H), 3.43 (s, 3H)

**13C NMR (100 MHz, CHLOROFORM-d):** δ 165.19, 163.98, 161.51 (d, J_CF_ = 246.9 Hz), 157.41, 155.34, 150.53, 141.43 (d, J_CF_ = 3.13 Hz), 131.41, 128.86 (d, J_CF_ = 8.54 Hz), 117.45 (d, J_CF_ = 22.6 Hz), 112.63, 109.71, 107.34, 98.86, 40.84

**HRMS (ESI) m/z:** calc’d for C_17_H_12_FNO_4_ [M+H^+^]: 314.0823 found: 314.0842

*7-(benzyl(methyl)amino)-2-oxo-2H-chromene-3-carboxylic acid **(7ACC2)***

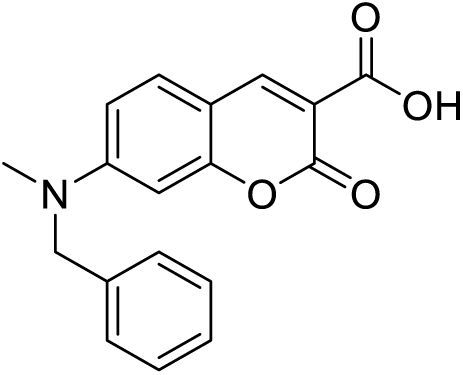

**1H NMR (500 MHz, CHLOROFORM-d):** δ 12.32 (s, 1H), 8.65 (s, 1H), 7.45 (d, J= 7.2 Hz, 1H), 7.38-7.27 (m, 3H), 7.17 (d, J= 6.8 Hz, 2H), 6.79 (dd, J= 1.20, 7.20 HZ, 1H), 6.60 (s, 1H), 4.72 (s, 2H), 3.25 (s, 3H).

**13C NMR (100 MHz, CHLOROFORM-d):** δ 165.36, 164.18, 157.66, 155.32, 150.47, 135.84, 131.82, 129.11, 127.83, 126.22, 111.35, 109.12, 106.58, 97.70, 56.31, 39.45.

**HRMS (ESI) m/z:** calc’d for C_18_H_15_NO_4_ [M+H^+^]: 310.1074 found: 310.1083

### Spectral characterization of 1,3-dihydroxy-2-(hydroxymethyl)propan-2-aminium 7-(methyl(4-(trifluoromethyl)benzyl)amino)-2-oxo-2H-chromene-3-carboxylate (D7)

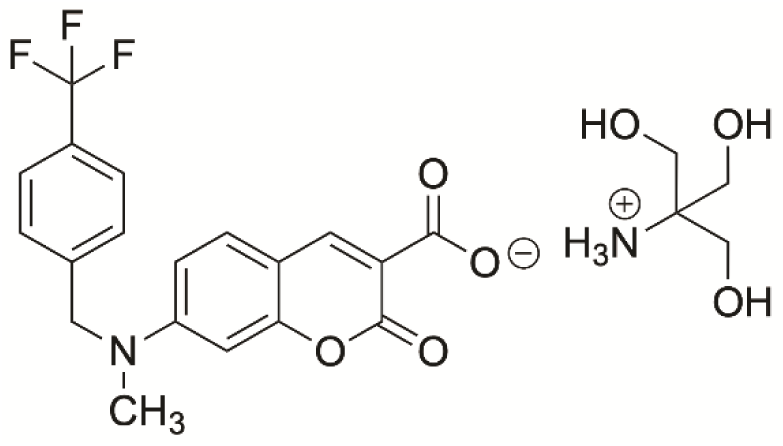

**^1^H NMR (400 MHz, DMSO-d6):** δ 8.209 (s, 1H), 7.6814 (d, 2H, J= 25.92), 7.5125 (d, 1H, J-8.72 Hz), 7.4255 (d, 2H, J= 7.68 Hz), 6.7399 (s, 1H, J= 8.8 Hz), 6.5566 (s, 1H), 4.8303 (s, 2H), 3.1767 (s, 3H)

**^13^C NMR (100 MHz, DMSO-d6):** δ 167.9491, 159.1321, 156.8593, 152.7093, 144.7865, 143.6035, 130.4881, 128.14085 (q, J= 31.5 Hz), 127.7736, 125.9397 (q, J= 3.6 Hz), 124.7483 (q, J= 270.4 Hz), 119.0451, 109.8017, 108.9669, 97.4376, 60.9012, 60.5096, 55.1307, 39.5101

**^19^F NMR ( 376 MHz DMSO-d6):** δ 60.8502

**HRMS (ESI) m/z:** calculated for C_19_H_14_F_3_NO_4_ [M+1Na]^+^: 400.0767, found 400.0774

## REFERENCES

1. Hanahan, D. and Weinberg, R. a (2000) The hallmarks of cancer. Cell 100, 57–70

2. Martinez-Outschoorn, U.E. et al. (2017) Cancer metabolism: a therapeutic perspective. Nat. Rev. Clin. Oncol. 14, 113–113

3. Albini, A. et al. (2015) Cancer stem cells and the tumor microenvironment: interplay in tumor heterogeneity. Connect. Tissue Res. 56, 414–425

4. Chang, C.H. et al. (2015) Metabolic Competition in the Tumor Microenvironment Is a Driver of Cancer Progression. Cell 162, 1229–1241

5. Hanahan, D. and Weinberg, R. A. (2011) Hallmarks of cancer: The next generation. Cell 144, 646–674

6. Curry, J.M. et al. (2013) Cancer metabolism, stemness and tumor recurrence : MCT1 and MCT4 are functional biomarkers of metabolic symbiosis in head and neck cancer. Cell Cycle 12, 1371–1384

7. Lee, M. (2015) Metabolic interplay between glycolysis and mitochondrial oxidation: The reverse Warburg effect and its therapeutic implication. World J. Biol. Chem. 6, 148

8. Fu, Y. et al. (2015) The reverse Warburg effect is likely to be an Achilles’ heel of cancer that can be exploited for cancer therapy. Oncotarget 5, 57813–57825

9. Ganapathy, V. et al. (2009) Nutrient transporters in cancer: Relevance to Warburg hypothesis and beyond. Pharmacol. Ther. 121, 29–40

10. Alvero, A.B. et al. (2014) Multiple blocks in the engagement of oxidative phosphorylation in putative ovarian cancer stem cells: implication for maintenance therapy with glycolysis inhibitors. Oncotarget 5, 8703–8715

11. Lee, N. and Kim, D. (2016) Cancer Metabolism: Fueling More than Just Growth. Mol. Cells 39, 847–854

12. Archetti, M. (2014) Evolutionary dynamics of the Warburg effect: Glycolysis as a collective action problem among cancer cells. J. Theor. Biol. 341, 1–8

13. Diaz-Ruiz, R. et al. (2011) The Warburg and Crabtree effects: On the origin of cancer cell energy metabolism and of yeast glucose repression. Biochim. Biophys. Acta - Bioenerg. 1807, 568–576

14. Lee, N. et al. (2016) Cancer Metabolism: Fueling More than Just Growth. Mol. Cells 39, 847–854

15. Xu, X.D. et al. (2015) Warburg effect or reverse warburg effect? a review of cancer metabolism. Oncol. Res. Treat. 38, 117–122

16. Bender, T. & Martinou, J. (2016) The mitochondrial pyruvate carrier in health and disease: To carry or not to carry? Mol. Cell Res. 1863, 2436–2442.

17. Tavoulari, S. et al. (2019) The yeast mitochondrial pyruvate carrier is a hetero-dimer in its functional state. 38, 1–13 doi:10.15252/embj.2018100785

18. Corbet, C. et al. (2018) Interruption of lactate uptake by inhibiting mitochondrial pyruvate transport unravels direct antitumor and radiosensitizing effects. Nat. Commun. 1–11. doi:10.1038/s41467-018-03525-0

19. Gurrapu, S. et al. (2015) Monocarboxylate transporter 1 inhibitors as potential anticancer agents. ACS Med. Chem. Lett. 6, 558–561.

20. Gurrapu, S. et al. (2016) Coumarin carboxylic acids as monocarboxylate transporter 1 inhibitors: In vitro and in vivo studies as potential anticancer agents. Bioorganic Med. Chem. Lett. 26, 3282–86.

21. Jonnalagadda, S., et al. (2019) Novel N, N-dialkyl cyanocinnamic acids as monocarboxylate transporter 1 and 4 inhibitors. 10, 2355–2368.

22. Draoui, N. et al. (2014) Antitumor Activity of 7-Aminocarboxycoumarin Derivatives, a New Class of Potent Inhibitors of Lactate Influx but Not Efflux. Mol. Cancer Ther. 13, 1410–1418.

